# Assessment of critically endangered Northern River Terrapin (*Batagur baska*) phylogeny through next-generation sequencing-based mitogenome analyses

**DOI:** 10.1101/2025.02.03.636247

**Authors:** Swati Nawani, Amirtha Balan, Asim Bashir, Vishnupriya Kolipakam, Abhijit Das, Samrat Mondol

**Affiliations:** Wildlife Institute of India, Dehradun, Uttarakhand 248001

**Keywords:** Freshwater turtle, genetics, Complete mitochondrial DNA, *Batagur baska*, Endangered species

## Abstract

The Northern River Terrapin, *Batagur baska* (Gray, 1830), is a critically endangered freshwater turtle, primarily found in the estuaries and tidal regions of large rivers in South and Southeast Asia. *B. baska* has experienced extensive population declines and extirpations across most of its range, with no confirmed wild populations remaining. To support conservation efforts of this turtle, our study utilizes first complete mitochondrial genome of *B. baska* using next-generation sequencing (NGS). This mitochondrial genome data is combined with existing genetic information to evaluate the species’ phylogenetic position within the Geoemydidae family. The Bayesian phylogenetic analysis, including 38 species sequences, confirmed a close genetic relationship between *B. baska* and B. *affinis* (posterior probability = 1). The mitochondrial genome of *B. baska* consists of 16,503 base pairs, with a typical structure comprising 13 protein-coding genes, two rRNA genes, and 22 tRNA genes, along with a non-coding control region. The protein-coding genes account for 68.9% of the genome, while the ribosomal RNA and tRNA genes cover 15.4%. The control region, located between tRNAPro and tRNAPhe, is 986 bp long and exhibits a distinct base composition.

## Introduction

The Northern River Terrapin, *Batagur baska*, (Gray, 1830) is one of the world’s most critically endangered fresh water turtle. It is an estuarine turtle, inhabiting across the estuaries and tidal portions of larger rivers. Historically, *B. baska* was reported from eastern India to Malaysia. Praschag et al. (2007) split this species based on morphology and genetic study into two species *B. baska* (Sundarband to Myanmar) and *B. affinis* (Indonesia, Cambodia, Malaysia, Thailand, and Vietnam). *B. baska* has suffered extensive extirpations across most of its former range, with no confirmed wild populations documented (Praschag et al., 2008) in recent times. During the 19^th^ century, Sundarban hosted a good population of *B. baska* but by 1970s the population reduced drastically (Moll, 1985), with only around 10 breeding females estimated by 1995 (Choudhury et al., 2000). This alarming decline has earned it a place among the 25 most critically endangered turtle species globally (Rhodin et al., 2011) and the classification of “ecologically extinct” (Weissenbacher et al., 2015). Recognizing its critical status CITES have included the species in Appendix I. The in-situ conservation program for *Batagur baska* was initiated in Sajnekhali, Sundarban Tiger Reserve (STR) in 2012 with an initial population of 12 individuals. Since the commencement of the program in 2012, the total population has increased to 397 individuals. The captive individuals are housed across seven camps, namely Sajnekhali, Dobanki, Jhila, Jhingekhali, Harikhali, Netidhopani, and Chamta. The critical need for *B. baska* conservation requires a multi-pronged approach encompassing both ecological and genetic studies. While ecological research has been the mainstay of its conservation efforts for decades, integrating genetic data offers a powerful complementary perspective (Corlett, 2016).

Thus, our study utilizes next-generation sequencing (NGS) to generate first complete mitochondrial genome data for *Batagur baska*. We then integrate this novel mitogenomic data with existing genetic information to comprehensively assess the phylogenetic position of this species within the Geoemydidae family.

## Materials and Methods

### Research permits

All required research permissions towards fieldwork and biological sampling were accorded by the Principal Chief Conservator of Forests (Wildlife) & Chief Wildlife Warden, West Bengal Forest Department (Permit Nos. 1304/WL/4R-28/2023 and 628/WL/4R-28/2023).

### Biological sampling

The biological sampling was conducted from the in-situ conservation program for *Batagur baska* situated in the Indian part of the Sundarbans mangrove forest, southern region of West Bengal, India. This region is situated on the Ganges Delta, formed by the confluence of the Ganges, Brahmaputra, and Meghna rivers, at the northern boundary of the Bay of Bengal (Iftekhar, 2006; Siddiqi, 2001), and is one of the places where this species was historically distributed (Moll, 1985, Choudhury et al., 2000). Approximately 40% of the Sundarbans (∼4,260 km^2^ area) lies within India, while the remaining (∼6,017 km^2^) is situated in Bangladesh (WCMC, 2005), characterized by a closed network of rivers, channels, and creeks, leading to the formation of numerous flat islands. This area is designated as an UNESCO World Heritage Site since 1987 and a Biosphere Reserve in 1989 (Ghosh et al., 2015).

During the 19^th^ century, Sundarbans hosted a good population of *B. baska* but this was drastically reduced by 1995 (Choudhury et al., 2000). The in-situ conservation program was initiated in Sajnekhali, Sundarbans Tiger Reserve (STR) in 2012 with an initial population of 12 individuals, resulting to the current population of ∼400 individuals spread across seven such camps, Sajnekhali (SJ), Dobanki DB), Jhila (JI), Jhingekhali (JH), Harikhali (HK), Netidhopani (ND), and Chamta (CM) **(Figure 1)**. During January 2024, blood sampling was conducted from four individuals during field visits to (Sajnekhali: n=2 and Dobanki: n=2) as per the standard procedures. Blood samples were collected aseptically (∼2 ml) by the Forest Department veterinary officials from the subcarapacial venous for each individual. The animals were released to their respective ponds after sampling. The collected blood was then preserved in EDTA-containing vacutainers (BD Diagnostics, USA) and transported to the laboratory at Wildlife Institute of India for storage at -20°C until further analysis.

**Figure 1:**
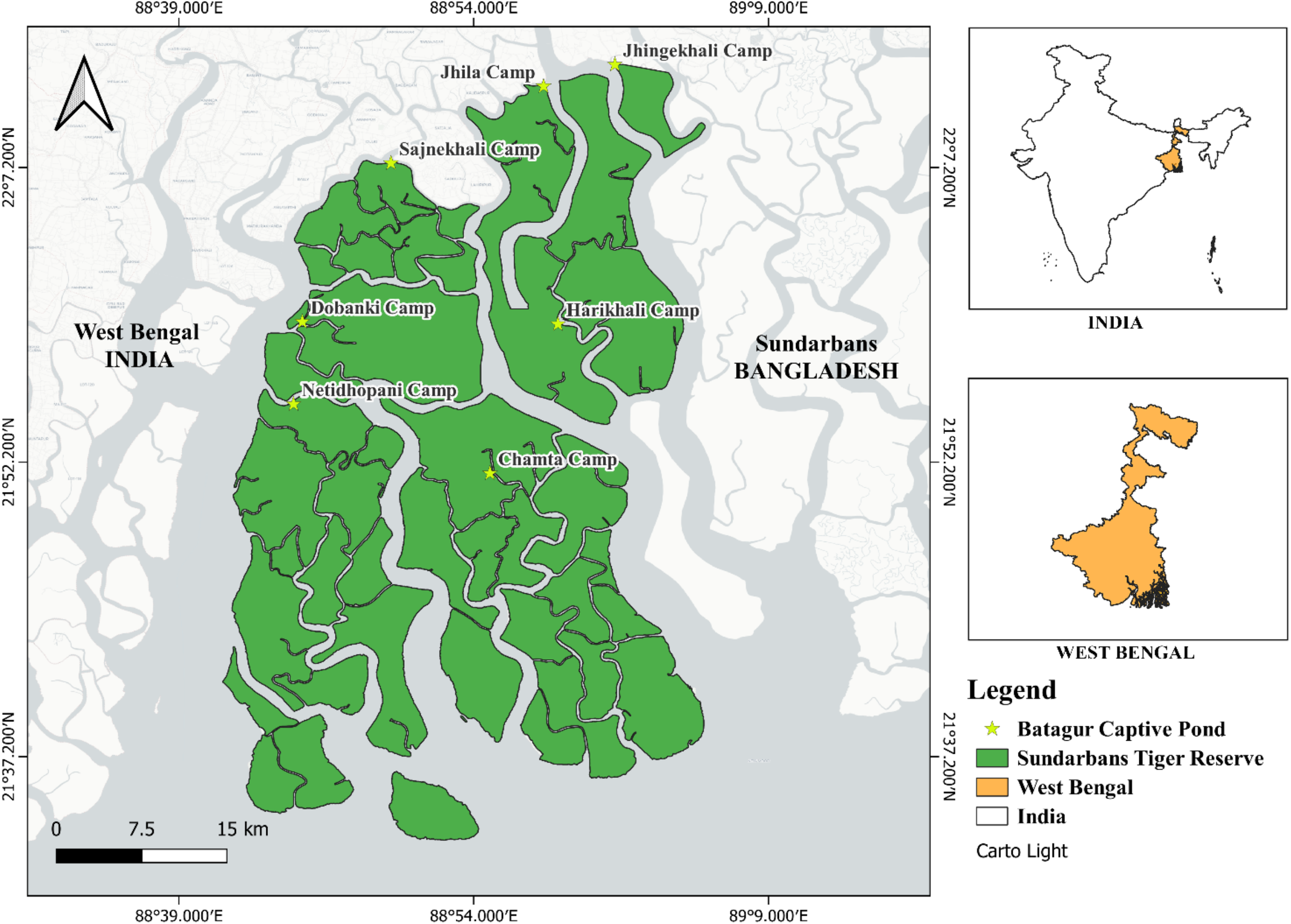
Sampling location in STR.

### DNA extraction and mitogenome sequencing

Mitogenome sequencing of *B. baska* was performed using next-generation sequencing (NGS) approach. Genomic DNA was extracted from all four blood samples using Alexgen Blood gDNA Kit (Alexius Biosciences, India) following manufacturers’ instructions. DNA quantity was checked through 1.8% agarose gel electrophoresis (LCO-1490, HCO2198 and 8F and 1492R primers) and Qubit 4.0 Fluorometer (Invitrogen, Life Technologies, USA). Subsequently, paired-end sequencing library was prepared using Twist NGS Library Preparation Kits () using 50 ng initial DNA inputs from each sample. The genomic DNA was enzymatically fragmented, end-repaired, A-tailed (at 3’ ends), and ligated with full length Illumina sequencing adapters at both ends. Following this, a high-fidelity amplification step was performed using HiFi PCR Master Mix. The amplified libraries were analyzed for purity, size, and concentration using TapeStation 4150 (Agilent Technologies, USA) using High Sensitivity D1000 ScreenTape® as per manufacturer’s instructions. Finally, the qualified libraries (2 × 150 bp) were loaded on Illumina NovaSeq6000 (Illumina, USA) platform for cluster generation and sequencing. The raw sequence reads were screened using Trimgalore 0.6.4 (http://www.bioinformatics.babraham.ac.uk/projects/trimgalore/) for quality assurances and remove the adapter sequences. After quality filtering, high-quality reads were assembled using Novoplasty 4.3.1 (Dierckxsens et al., 2017), CLC Genomics Workbench 24 (Matvienko, 2015), and animal mtDNA database sequences integrated in GetOrganelle ver 1.7.7.0 (Jin et al., 2020)

All *B. baska* sequences (n=4) were checked manually and cleaned for any nucleotide ambiguities and the overlapping regions were aligned to generate a complete mitogenome sequence using Mega v7 (Kumar et al., 2016). The resulting sequence was then annotated using the MITOS2 online server (http://mitos.bioinf.uni-leipzig.de). The circular representation of the *B. baska* mitogenome was generated using the OGDRAW Server (https://chlorobox.mpimp-golm.mpg.de/OGDraw.html) with default parameters. Intergenic spacers and overlapping regions were manually counted using Microsoft Excel. Additionally, nucleotide percentages were calculated using Mega v7. Skew analysis was conducted using the following formulas: GC skew = (G-C)/(G+C); AT skew = (A-T)/(A+T) to evaluate nucleotide composition bias.

### Phylogenetic analysis

For phylogenetic analyses, a total of 38 Geoemydidae family mitogenome sequences were downloaded from Genbank. These sequences were aligned with the *B. buska* mitogenome generated in this study, and the mean pairwise genetic distances among them was estimated using Mega v7 (Kumar et al., 2016). Two sequences of *Chitra indica* and *Lissemys punctata* were used as outgroup in this analysis.

A two-parameter substitution model (GTR+G, decided using jModelTest v. 2.1.10 (Darriba et al., 2016)) along with a gamma distribution of evolutionary rates (across sites), and default shape parameter setting were used in the phylogenetic analyses through BEAST v.1.6.1 (Drummond & Rambaut, 2007).). The MCMC parameters incorporated 100,000,000 generations and default settings. The phylogenetic tree was constituted with the protein-coding genes (PCG). No molecular clock was applied in this analysis, focusing on topology rather than distance for tree and branch length representation. The trees were visualized using FigTree (v. 1.4.4; http://tree.bio.ed.ac.uk/software/figtree/)

## Results

### Mitogenome structure and organization

A total of 22,7374,62 **(SJ1)**, 22,112,184 **(SJ2)**, 25,786,510 **(DB3)** and 25,355,932 **(DB4)** sequence reads were generated through NGS approach. The assembled mitogenome of *B. baska* generated in this study was of 16,503 bp length and matched with the available *Batagur* species reference sequences. The annotated *B. baska* mitogenome showed conserved gene order like other *Batagur* species i.e. protein-coding genes (PCGs, n = 13), tRNA genes (n = 22), ribosomal genes (n = 2) and a non-coding control region (Table 1). Out of 37 genes, NADH6 and eight tRNA genes (tRNAGln, tRNAAsn, tRNAAla, tRNACys, tRNATyr, tRNASer, tRNAGlu and tRNAPro) were encoded on the light strand (L-strand), whereas all others (n=29) are encoded on the heavy strand (H-strand). The nucleotide composition of *B. baska* mitogenome consisted of 24.3% T, 28.8% C, 33.7% A and 13.2% G. Composition analyses through AT and GC skew showed a positive AT value (0.16) and a negative GC value (−0.37), indicating a AT-rich mitogenome composition of Northern River Terrapin.

**Table 1:** Organization of mitochondrial genome in *B. baska*.

| Gene | Start | End | Spaces/Overlap | Start codon | Stop codon | Strand |
| --- | --- | --- | --- | --- | --- | --- |
| trnF(gaa) | 1 | 69 | 0 |  |  | + |
| rrnS | 70 | 1033 | 0 |  |  | + |
| trnV(tac) | 1034 | 1103 | 12 |  |  | + |
| rrnL | 1115 | 2699 | 2 |  |  | + |
| trnL2(taa) | 2701 | 2776 | 0 |  |  | + |
| nad1 | 2777 | 3745 | -1 | ATG | TAG | + |
| trnI(gat) | 3745 | 3814 | -1 |  |  | + |
| trnQ(ttg) | 3814 | 3884 | -1 |  |  | - |
| trnM(cat) | 3884 | 3952 | 0 |  |  | + |
| nad2 | 3953 | 4993 | -2 | ATG | TAG | + |
| trnW(tca) | 4992 | 5062 | 2 |  |  | + |
| trnA(tgc) | 5064 | 5132 | 2 |  |  | - |
| trnN(gtt) | 5134 | 5206 | 27 |  |  | - |
| trnC(gca) | 5233 | 5298 | 8 |  |  | - |
| trnY(gta) | 5306 | 5376 | 2 |  |  | - |
| cox1 | 5378 | 6928 | -11 | GTG | TAA | + |
| trnS2(tga) | 6917 | 6987 | 0 |  |  | - |
| trnD(gtc) | 6988 | 7057 | 0 |  |  | + |
| cox2 | 7058 | 7744 | 2 | ATG | TAA | + |
| trnK(ttt) | 7746 | 7818 | 2 |  |  | + |
| atp8 | 7820 | 7987 | -9 | ATG | TAA | + |
| atp6 | 7978 | 8661 | -1 | ATG | TAA | + |
| cox3 | 8661 | 9445 | -1 | ATG | TAA | + |
| trnG(tcc) | 9445 | 9512 | 2 |  |  | + |
| nad3_1 | 9514 | 9693 | -4 | ATG | TAA | + |
| nad3_0 | 9689 | 9862 | 2 | ATG | TAA | + |
| trnR(tcg) | 9864 | 9933 | 0 |  |  | + |
| nad4l | 9934 | 10230 | -6 | ATG | TAA | + |
| nad4 | 10224 | 11600 | 19 | ATG | TAA | + |

**Table 1:** Organization of mitochondrial genome in *B. baska*
|  |  |  |  |  |  |  |
| --- | --- | --- | --- | --- | --- | --- |
| trnH(gtg) | 11619 | 11687 | 0 |  |  | + |
| trnS1(gct) | 11688 | 11756 | -1 |  |  | + |
| trnL1(tag) | 11756 | 11828 | 0 |  |  | + |
| nad5 | 11829 | 13637 | -4 | ATG | TAA | + |
| nad6 | 13633 | 14157 | 0 | ATG | TAA | - |
| trnE(ttc) | 14158 | 14225 | 6 |  |  | - |
| cob | 14231 | 15370 | 5 | ATG | AGA | + |
| trnT(tgt) | 15375 | 15447 | 0 |  |  | + |
| trnP(tgg) | 15448 | 15518 | 0 |  |  | - |
| D loop | 15519 | 16503 | - |  |  | + |

### Protein Coding Genes

The total length of the Northern River terrapin mitochondrial protein coding genes was 11369 bp, accounting about 68.9% of the mitogenome. Average base composition was 24.5% T, 30.8% C, 32.8% A and 11.9% G suggesting that AT% was more than GC% in the protein coding region. The coding genes included seven NADH genes (NADH1, NADH2, NADH3, NADH4, NADH5, NADH6, and NADH4L), two ATPase (ATP6 and ATP8), three cytochrome c oxidases (COI, COII and COIII) and one cytochrome b (Cyt b). These protein coding regions were AT skewed. Majority of these genes (n=12) were present in the H-strand. All genes started with ATG except COI which starts with GTG.

### Ribosomal RNA (rRNA) and transfer RNA (tRNA) genes

Northern River Terrapin mitogenome has two rRNA genes: 12s rRNA and 16s rRNA, that are located between tRNAPhe and tRNAVal, and tRNAVal and tRNALeu, respectively (Figure 3). Total sequence length of ribosomal RNA was 2549 bp. The mitogenome also contains 22 tRNA genes covering a total length of 1547 bp. Average base composition was 35% A, 24.4% T, 15.9% G and 24.8% C. Their length varied from 65bp (tRNAc) to 75 bp (tRNAL2). Out of the 22 genes, 14 were located on H-strand and rest eight were found on L-strand.

### Control region

The Control region (CR) of *B. baska* is 986 bp long and located between tRNAPro and tRNAPhe. The base composition of CR was 31.8% T, 22.1% C, 32% A and 14% G. **Phylogenetic Relationship**

Mitogenome-based Bayesian phylogenetic reconstruction of the *B. baska* (with 38 available species sequences including outgroups) confirmed its relationship with *B. affinis* with high posterior probability (PP = 1) **(Figure 2)**. The obtained Testudines phylogeny and the relationship of *Batagur* are largely consistent with previous results (Kundu et al., 2019; Kumar et al., 2021). Additionally, the genus *Cuora* is close to *Mauremys*, whereas *Cyclemys, Sacalia, Heosemys*, and *Notochelys* are clustered separately with high posterior probability support.

**Figure 2:**
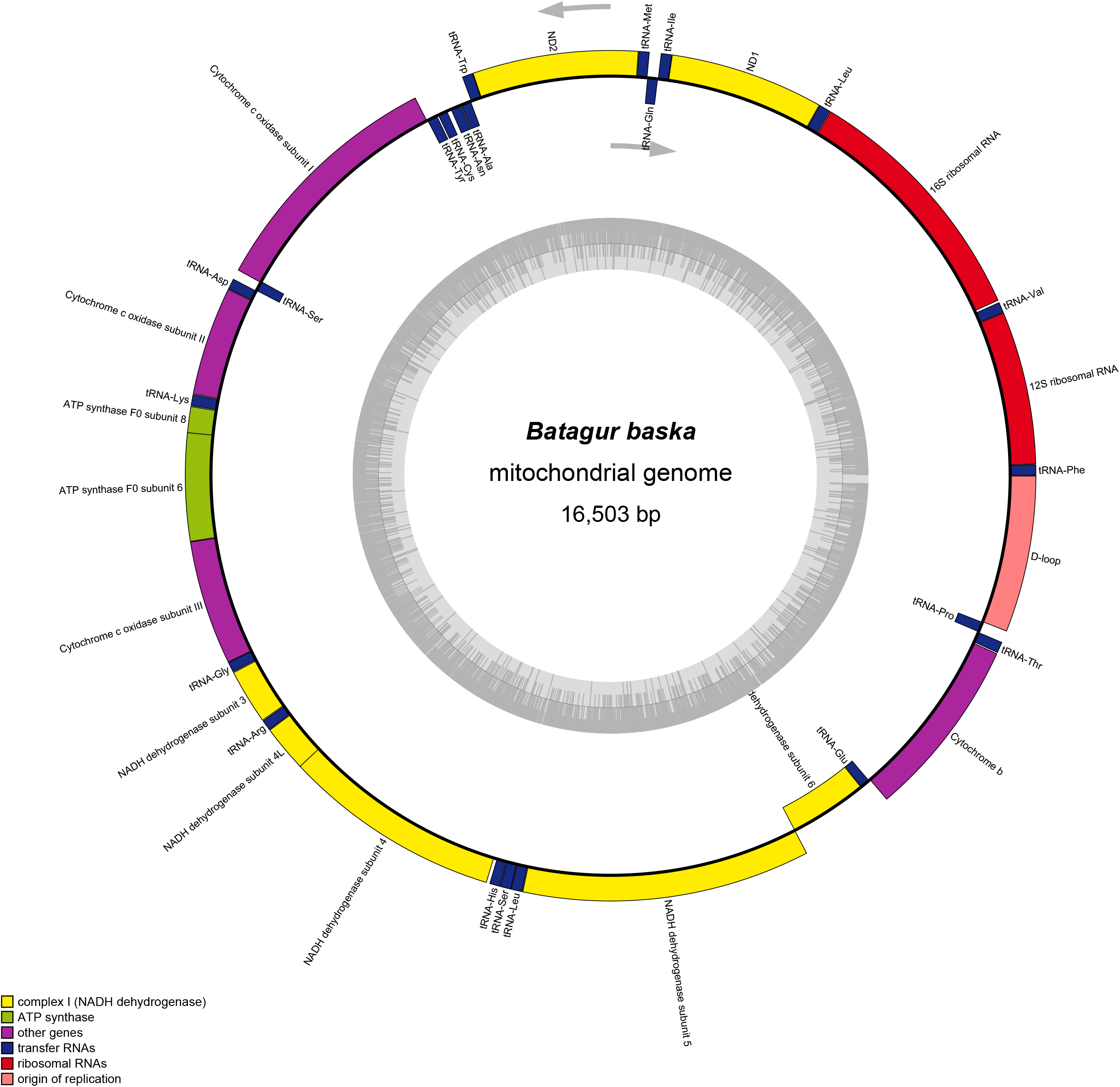
Whole mitogenome map of *Batagur baska*.

**Figure 3:**
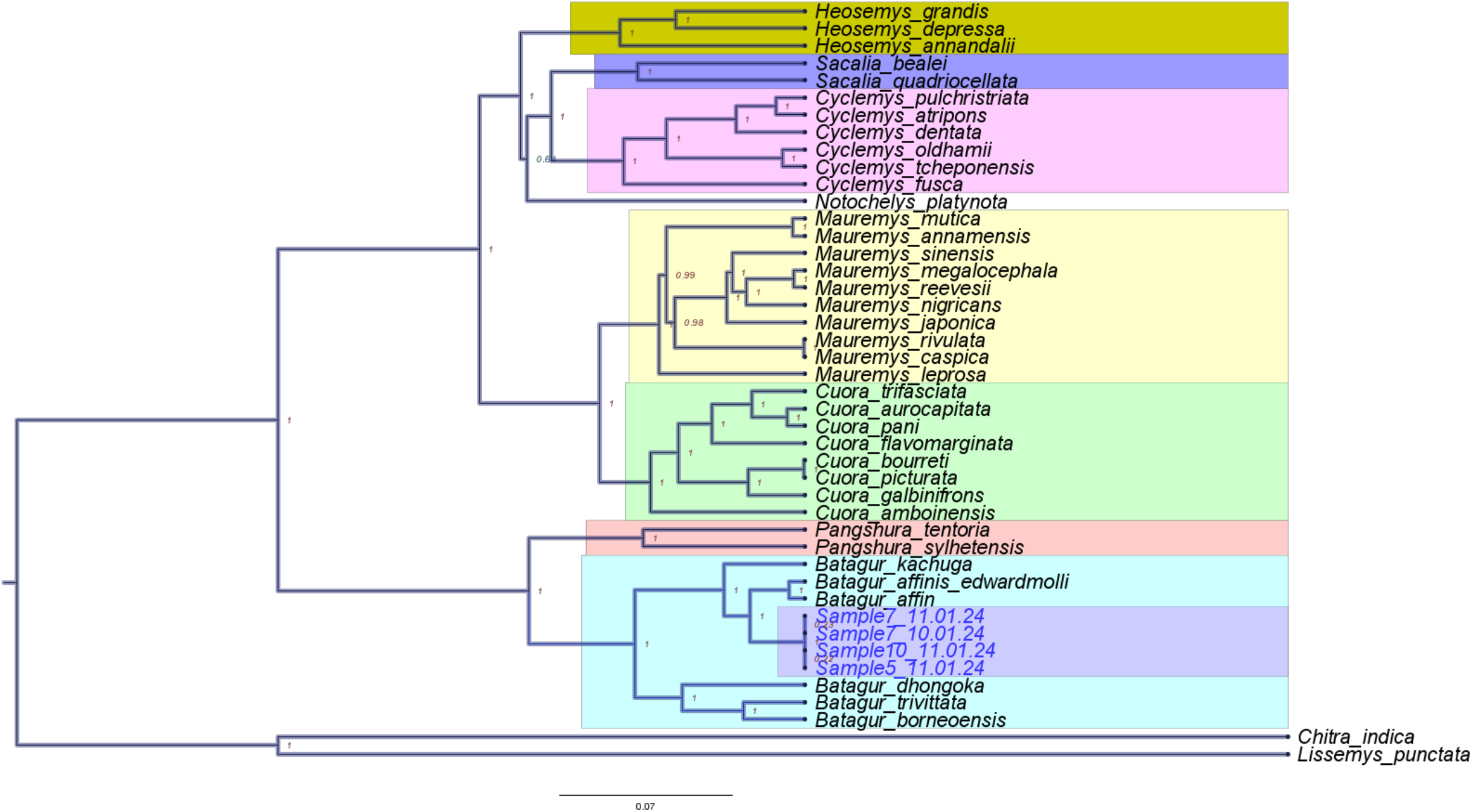
Phylogenetic relationship of *Batagur baska* inferred from Bayesian inference analysis using 13 concatenated protein coding genes. Posterior probability values are shown at the nodes of the tree.

## Discussion

This study has provided valuable insights into whole mitochondrial genome of *Batagur baska*. A recent study by Das et al. (2024) determined the complete mitogenome sequence of *Batagur kachuga* which possesses a 16,517 bp mitogenome. This size is remarkably similar to that of *B. baska*. However, a previous study by Kumar et al. (2021) reported a smaller mitogenome size of 16,155 bp for *B. baska*. Comparative analysis with other *Batagur* species reveals interesting patterns. Kumar et al. (2021) reported a smaller mitogenome size of 15,619 bp for *Batagur dhongoka*. In contrast, Feng et al. (2017) estimated a mitogenome size of 16,463 bp for *Batagur trivittata*. The *B. baska* mitogenome exhibits a conserved gene order with other *Batagur* species characterized by an AT-rich bias aligning with observations in other turtles. The comparative analysis of mitogenome structures within the *Batagur* genus reveals a high degree of conservation. *B. baska, B. kachuga, B. dhongoka*, and *B. trivittata* share a typical vertebrate mitogenome organization, comprising 13 protein-coding genes, 22 transfer RNA genes, two ribosomal RNA genes (12S rRNA and 16S rRNA), and a non-coding control region. A notable feature is the strand asymmetry, with the majority of genes (28-29) encoded on the heavy strand (H-strand) and a smaller number (8-9 genes) on the light strand (L-strand). This strand bias is a common characteristic of vertebrate mitogenomes and is likely related to replication and transcription efficiency (Fonseca et al., 2014; Gomes-dos-Santos et al., 2023). Thus, while the overall gene content and arrangement are highly conserved across these species, minor variations exist.

Comparative mitogenomic data have also demonstrated utility in elucidating the phylogenetic relationships of turtles (Kundu et al., 2019). The present mitogenomes study was able to generate a robust phylogeny with high posterior probability value for each node and elucidate the relationship between *B. baska* and other Geoemydid species.

*B. baska*, historical records indicate its local extirpation in Orissa, India (Behera et al., 2019), and its population status in Myanmar remains uncertain (Moll et al., 2009) suggesting a dire need of further studies on its genetic makeup and ecology. Sundarbans, once a thriving habitat, is increasingly degraded due to human activities like deforestation, pollution, and unsustainable fishing practices. Additionally, climate change-induced sea-level rise pose additional threats. Hence, a deeper understanding of the genetic diversity and population structure of *B. baska* will aid in identifying genetically distinct populations, assessing the impact of habitat fragmentation, and prioritizing conservation efforts. By analyzing genetic markers, we can identify potential threats such as inbreeding depression and genetic drift, which can compromise the long-term viability of populations. Furthermore, genetic information can be used to inform captive breeding programs and reintroduction efforts, ensuring the health of future generations.

## Acknowledgment

We acknowledge the Principal Chief Conservator of Forests (Wildlife) & Chief Wildlife Warden, West Bengal Forest Department for research permits 1304/WL/4R-28/2023 dated 17.05.2023 and **628/WL/4R-28/2023 dated 10/3/2023** and logistic support during fieldwork. Our sincere thanks to the Field Director and Deputy Field Director and all the staff’s at Sundarban Tiger Reserve for providing necessary assistance during our fieldwork. We are indebted to forest departments Rangers Mr. Biplab Ghosh, Mr. Avik Das, Mr. Sumith Bose, Mr. Swapan Majhi, Mr. Saeef and other Staffs Mr. Sabir Ansari, Mr. Nabukumar, Mr. Gautam Dali, Mr. Monirudhin. During fieldwork support was provided by Mr. Tarun Das, Mr. Ardhendu Mondal and Mr. Prodip Mirdha for their constant support and care during our field work.

This work was possible through the financial assistance from Ministry of Environment, Forest and Climate Change, Govt of India under its IDWH scheme. Our sincere thanks to the Director, Registrar, Dean, Research Coordinator, Nodal Officer WFCG Cell, Laboratory in-charge, Special thanks to Dr. K. Sivakumar of Pondicherry university for his support during formulation of the project. We thank Dr. Gowri Mallapur and Dr. Anukul Nath for their kind support and guidance. At WII, Mr. Ankit Pacha, Miss Shrewshree Kumar, and Miss Megha Mehta for their assistance in data analyses.

